# Chromomycin A_2_ potently inhibits glucose-stimulated insulin secretion from pancreatic β cells

**DOI:** 10.1101/337113

**Authors:** Michael A Kalwat, In Hyun Hwang, Jocelyn Macho, Magdalena G Grzemska, Jonathan Z Yang, Kathleen McGlynn, John B MacMillan, Melanie H Cobb

## Abstract

Enhancers or inhibitors of insulin secretion could become therapeutics as well as lead to the identification of requisite β-cell regulatory pathways and increase our understanding of pancreatic islet function. Toward this goal, we previously used an insulin-linked luciferase that is co-secreted with insulin in MIN6 β-cells to perform a high-throughput natural product screen for chronic effects on glucose-stimulated insulin secretion. Using multiple phenotypic analyses, we identified that one of the top natural product hits, chromomycin A2 (CMA2), potently inhibited insulin secretion through at least three mechanisms: disruption of Wnt signaling, interfering with β-cell gene expression, and suppression of triggering calcium (Ca^2+^) influx. Chronic treatment with CMA2 largely ablated glucose-stimulated insulin secretion even post-washout, but did not inhibit glucose-stimulated generation of ATP or Ca^2+^ influx. However, by using the K_ATP_ channel-opener diazoxide, we uncovered defects in depolarization-induced Ca^2+^ influx which may contribute to the suppressed secretory response. Glucose-responsive ERK1/2 and S6 phosphorylation were also disrupted by chronic CMA2 treatment. The FUSION bioinformatic database indicated that the phenotypic effects of CMA2 clustered with a number of Wnt/GSK3 pathway-related genes. Consistently, CMA2 decreased GSK3 phosphorylation and suppressed activation of a β-catenin activity reporter. CMA2 and a related compound mithramycin are described to have DNA-interaction properties, possibly abrogating transcription factor binding to critical β-cell gene promoters. We observed that CMA2, but not mithramycin, suppressed expression of PDX1 and UCN3. However, neither expression of INSI/II nor insulin content was affected by chronic CMA2. The mechanisms of CMA2-induced insulin secretion defects may involve components both proximal and distal to Ca^2+^ influx. Therefore, CMA2 is an example of a chemical that can simultaneously disrupt β-cell function through both non-cytotoxic and cytotoxic mechanisms. Future applications of CMA2 and similar aureolic acid analogs for disease therapies should consider the potential impacts on pancreatic islet function.

## Materials and Methods

### Antibodies and Reagents

Chromomycin A2 was purified from extracts of *Streptomyces anulatus* as described below or purchased (Santa Cruz). Mithramycin was generously provided by Ralf Kittler (UT Southwestern). Antibodies, their dilutions for immunoblotting, and their sources are listed in **Table 1**. All other reagents were obtained through Fisher unless otherwise stated.

### Isolation of natural product fractions and bacterial strain information

The library of microbial and invertebrate natural product fractions was subjected to liquid chromatography-mass spectrometry (LC-MS) analysis using an Agilent Model 6130 single quadrupole mass spectrometer with an HP1200 HPLC. A photodiode array detector provided a chemical fingerprint of all fractions on the basis of molecular weight and UV profile. Fractions were dereplicated using various compound databases, including Antibase, Reaxys, and Scifinder Scholar. The sediment was desiccated and stamped onto acidic agar plates (20 g starch, 1 g NaNO_3_, 0.5 g K_2_HPO_4_, 0.5 g MgSO_4_, 0.5 g NaCl, 0.01 g FeSO_4_, 10 μM cadaverine, 10 μM spermidine, 750 mL seawater, 15 g agar, pH adjusted to 5.3 with phosphate buffer). Bacterial colonies were selected and streaked for purity using the same agar medium. Standard procedures for 16S rRNA analysis were used for phylogenetic characterization of bacterial strains. Analysis of the *Streptomyces sp.* strain SN-B-022-1 by 16S rRNA revealed 99% identity to *Streptomyces anulatus.*

### Cultivation and Extraction of SN-B-022-1

Bacterium SN-B-022 was cultured in 5 × 2.8 L Fernbach flasks each containing 1 L of a seawater-based medium (10 g starch, 4 g yeast extract, 2 g peptone, 1 g CaCO_3_, 40 mg Fe_2_(SO_4_)_3_·4H_2_O, 100 mg KBr) and shaken at 200 rpm at 27°C. After 7 days of cultivation, sterilized XAD-7-HP resin (20 g/L) was added to adsorb organic chemicals produced by the bacterium, and the culture and resin were shaken at 200 rpm for 2 h. The resin was extracted with acetone and filtered through cheesecloth. The acetone-soluble portion was dried *in vacuo* to yield 2.0 g of extract.

### Purification of chromomycin A2 from SN-B-022-1

The extract (300 mg out of 2.0 g) was subjected to reversed-phase medium-pressure liquid chromatography (MPLC) with 5% CH_3_OH/H_2_O for 1 min, followed by a linear gradient of CH_3_OH/H_2_O from 5% to 100% over 15 min, 4 min of column washing with pure CH_3_OH, and substitution of the column with 5% CH_3_OH/H_2_O for 2 min to provide 20 library fractions (SNB-022-1 to - 20). Chromomycin A2 (1.2 mg, tR = 32.1 min) was obtained from the active fraction SN-B-022-1 (88 mg) by reversed-phase HPLC (5-μm, 10.0 mm × 250 mm) using 25% CH_3_OH/H_2_O (0.1% formic acid) for 5 min, followed by a linear gradient of CH_3_OH/H_2_O from 25% to 85% over 30 min, and 5 min of column washing with pure CH3CN. Application of the same HPLC purification method to SN-B-022-3 (14 mg) enabled isolation of additional samples of chromomycin A2 (2.6 mg). The compound was identified by comparison of NMR data to literature values (Toume, K. et al., 2014), and the structure of chromomycin A2 was confirmed independently through analysis of two-dimensional NMR (HSQC and HMBC) data.

### Cell culture and treatments

MIN6 β cells (originally obtained from John Hutton, University of Colorado) were cultured in Dulbecco’s modified Eagle’s medium (which contains 25 mM glucose), supplemented with 15% fetal bovine serum, 100 units/ml penicillin, 100 μg/ml streptomycin, 292 μg/ml L-glutamine, and 50 μM β-mercaptoethanol as described previously (Kalwat, M.A. et al., 2016a). MIN6 cells were seeded in 12-well dishes at ~4e5 cells per well and grown 4-7 d with medium changed every 2-3 d before use in experiments. For CMA2 or mithramycin treatments for RNA isolation, cells were treated overnight and harvested after 24 h. Prior to glucose stimulation experiments, MIN6 cells were washed twice with and incubated for 2 h in freshly prepared glucose-free modified Krebs-Ringer bicarbonate buffer (KRBH: 5 mM KCl, 120 mM NaCl, 15 mM HEPES, pH 7.4, 24 mM NaHCO3, 1 mM MgCl_2_, 2 mM CaCl_2_, and 1 mg/ml radioimmunoassay-grade BSA). Cells were lysed in 25 mM HEPES, pH 7.4, 1% Nonidet P-40, 10% glycerol, 50 mM sodium fluoride, 10 mM sodium pyrophosphate, 137 mM NaCl, 1 mM sodium vanadate, 1 mM phenylmethylsulfonyl fluoride, 10 μg/ml aprotinin, 1 μg/ml pepstatin, 5 μg/ml leupeptin and cleared of insoluble material by centrifugation at 10,000 x g for 10 min at 4°C for subsequent use. Human islets and MIN6 cells were stimulated with glucose as indicated and secreted insulin and insulin content were measured using ELISA (Mercodia) and HTRF (Cisbio) assays. For β-catenin reporter assays, stable Super TOPFlash HEK293 cells were plated 1e4/well and cultured for 48 h prior to treatment with indicated doses of CMA2 or CHIR99021 for 24 h. Medium was decanted and 20 μl of passive lysis buffer was added. After a 10 min incubation, 80 μl of Promega LARII firefly luciferase substrate working solution was added with an injector on a Synergy2 H1 plate reader (BioTek). Protein concentration of HEK293 cell lysates was measured by Bradford assay and SuperTOPFlash signal was normalized to the protein concentration.

### Human islet culture and treatments

Cadaveric human islets were obtained through the Integrated Islet Distribution Program (IIDP). Islets were isolated by the affiliated islet isolation center and cultured in PIM medium (PIM-R001GMP, Prodo Labs) supplemented with glutamine/glutathione (PIM-G001GMP, Prodo Labs), 5% Human AB serum (100512, Gemini Bio Products), and ciprofloxacin (61-277RG, Cellgro, Inc) at 37°C and 5% CO_2_ until shipping at 4°C overnight. Details of Donor information is provided in **Table 2**. Upon receipt, human islets were cultured in CMRL-1066 containing 10% fetal bovine serum, 100 U/ml penicillin and 100 μg/ml streptomycin. For static culture insulin secretion experiments, human islets were hand-picked under a dissection microscope equipped with a green Kodak Wratten #58 filter (Finke, E.H. et al., 1979) and placed into low-binding 1.5 ml tubes with ~50 islets per tube. Islets were washed twice with Krebs-Ringer Bicarbonate Hepes buffer (KRBH) (134 mM NaCl, 4.8 mM KCl, 1 mM CaCl_2_, 1.2 mM MgSO_4_, 1.2 mM KH_2_PO_4_, 5 mM NaHCO_3_, 10 mM HEPES pH 7.4, 0.1% BSA) and preincubated in KRBH supplemented with 2 mM glucose for 1 h before switching to KRBH containing either 2 or 16 mM glucose for 1 h. Supernatants were collected, centrifuged at 10,000 x g for 5 min and transferred to fresh tubes for storage at −80°C. Total insulin content was extracted by acid-ethanol overnight at −80°C and was neutralized with an equal volume of 1 M Tris pH 7.4 prior to assay.

### Secreted Gaussia luciferase assays

InsGLuc-MIN6 cells were generated as previously described (Kalwat, M.A. et al., 2016b). Cells were plated in 96-well dishes at 1e5 cells/well and incubated for 2-3 days before overnight (24 h) treatment with natural products. Cells were then washed twice with KRBH and preincubated in 100 μl of KRBH (250 μM diazoxide included where indicated in figure legends) for 1 h. The solution was then removed and cells were washed one time with 100 μl of KRBH and then incubated in KRBH with or without the indicated glucose concentration (or 250 μM diazoxide, 35 mM KCl, 20 mM glucose where indicated) for 1 h. 50 μl of KRBH was collected from each well and pipetted into a white opaque 96-well plate Optiplate-96 (Perkin-Elmer). Fresh Gaussia luciferase assay working solution was then prepared by adding coelenterazine stock solution into assay buffer to a final concentration of 10 μM. 50 μl of working solution was then rapidly added to the wells using a Matrix 1250 μl electric multi-channel pipette for a final concentration of 5 μM coelenterazine. After adding reagent to the plate, plates were spun briefly and luminescence was measured on a Synergy2 H1 plate reader (BioTek). The protocol was set to shake the plate orbitally for 3 seconds and then read the luminescence of each well with a 100 ms integration time and gain set to 150.

### Immunoblotting and Microscopy

40-50 μg of cleared cell lysates were separated on 10% gels by SDS-PAGE and transferred to nitrocellulose for immunoblotting. All membranes were blocked in Odyssey blocking buffer (Licor) for 1 h before overnight incubation with primary antibodies diluted in blocking buffer. After three 10 min washes in 20 mM Tris-HCl pH 7.6, 150 mM NaCl, 0.1% Tween-20 (TBS-T), membranes were incubated with fluorescent secondary antibodies for 1 h at room temperature. After three 10 min washes in TBS-T, membranes were imaged on a Licor Odyssey scanner. For experiments in which cells were fixed and stained with DAPI, MIN6 cells were plated in 96 well plates at 5e4 cells/well 24 h prior to treatment. After treatment, cells were fixed and permeabilized in 4% paraformaldehyde and 0.18% Triton-X100 in PBS pH 7.4 for 10 min, washed three times for 5 min each, and then stained with DAPI (300 nM) in PBS for 10 min. The cells were washed again twice with PBS and taken immediately for imaging. Plates were imaged in the UT Southwestern High-throughput Screening core facility using a GE^®^INCell 6000 automated microscope (GE Healthcare), 10X objective lens, at room temperature, and using a DAPI filter. Nuclei counting was performed using a CellProfiler workflow (Carpenter, A.E. et al., 2006).

### Quantitative PCR (RT-qPCR)

RNA was isolated from MIN6 cells using the PureLink RNA Mini Kit (Life Technologies). 500 ng of MIN6 RNA was converted into cDNA using the iScript kit (Bio-Rad) and the resulting cDNA was diluted 10-fold with water. One μl of diluted cDNA was used in 10 μl qPCR reactions using 2X SYBR Bio-Rad master mix and 250 nM of each primer. qPCR data was analyzed using QuantStudio software (Applied Biosystems) and 18S RNA was used as the reference gene. All primer sequences are listed in **Table 3**.

### Statistical Analysis

Quantitated data are expressed as mean ± SE. Data were evaluated using unpaired Student’s t test or two-way ANOVA where indicated, and considered significant if P < 0.05. Graphs were made in GraphPad Prism 6.

## INTRODUCTION

Insulin secretion from β cells is one of the most important functions of the pancreatic islet. Type 1 and type 2 diabetes combined afflict 9.4% of Americans and result from the autoimmune destruction of β cells, or defects in insulin secretion and action, respectively. Diabetes is usually treated by insulin replacement or by β cell-targeted therapeutics which are typically inducers of insulin secretion. However, insulin hypersecretion also occurs in type 2 diabetes and other disorders such as persistent hyperinsulinemic hypoglycemia of infancy and polycystic ovary syndrome (Stanley, C.A., 2011, Molven, A. et al., 2003, Cordain, L. et al., 2003). Accordingly, suppression of insulin secretion has been considered in the past as a therapeutic avenue (Hansen, J. et al., 2004). Pancreatic islet β cells sense hyperglycemia and respond by metabolizing glucose, increasing their intracellular [ATP/ADP] ratio and causing the closure of ATP-sensitive potassium (K_ATP_) channels. These steps lead to the opening of voltage-dependent Ca^2+^ channels, Ca^2+^ influx and insulin granule exocytosis denoted as ‘triggering’ (Henquin, J.C., 2000). Glucose metabolism also mediates an ‘amplifying’ pathway through the production of other signals that drive insulin secretion independent from further Ca^2+^ influx (Gembal, M. et al., 1992, Sato, Y. et al., 1992). Much is known about the triggering pathway; however, less is understood about the glucose-mediated metabolic amplifying pathway which can account for half of the insulin secretion response (Henquin, J.C. et al., 2017, Henquin, J.C., 2009). In particular, the signaling pathways and transcriptional outputs that are chronically required for regulated insulin secretion are not completely understood. Further development of knowledge and tools pertaining to these aspects of β cell function are required to uncover new directions for research and to guide disease therapies.

Marine natural products are a rich resource for potential new chemical tools and therapies. A unique custom library of natural compounds assembled at UT Southwestern has been utilized in screens to discover new leads for chemotherapies (Hu, Y. et al., 2011, Vaden, R.M. et al., 2017, Potts, M.B. et al., 2015, Potts, M.B. et al., 2013), inhibitors of endocytosis (Elkin, S.R. et al., 2016), antibiotics (Hu, Y. et al., 2013), and, in our recent screening efforts, modulators of pancreatic β cell function (Kalwat, M.A. et al., 2016b). As a result, we discovered that the natural product chromomycin A_2_ (CMA2) is a potent inhibitor of insulin secretion. CMA2 is a member of a family of glycosylated aromatic polyketides called aureolic acids. This family includes a variety of chromomycins, olivomycins, chromocyclomycin, duhramycin A, as well as the founding member mithramycin (Lombo, F. et al., 2006). This class of compounds is known to interact with the minor groove of DNA in a non-intercalative way with a preference for G/C-rich regions (Hou, C. et al., 2016). This mechanism is more subtle and distinct from other natural product transcription inhibitors like actinomycin D which binds DNA at the transcription elongation complex (Sobell, H.M., 1985) or α-amanatin which directly inhibits RNA polymerase II (Bushnell, D.A. et al., 2002). Based on their known activities, aureolic acids have been pursued as anti-cancer chemotherapies. In addition to their anti-cancer properties (Lombo, F. et al., 2006, Singh, D.K. et al., 2017), these compounds have been implicated as inhibitors of Wnt signaling (Toume, K. et al., 2014) and DNA gyrase (Simon, H. et al., 1994), as well as inducers of autophagy (Guimaraes, L.A. et al., 2014, Ratovitski, E.A., 2016). Despite this array of activities, the specific genes and pathways involved in the mechanisms of action of CMA2 and mithramycin are only just being fleshed out. Here we have determined that CMA2 alters β-cell function by interfering with Ca^2+^ influx, suppressing β cell gene expression, and disrupting GSK3/β-catenin signaling.

## RESULTS

### Identification of Chromomycin A_2_ as an insulin secretion inhibitor

We previously performed a natural product screen using specialized MIN6 β cells containing an insulin promoter-driven Gaussia luciferase (GLuc)-linked insulin transgene, InsGLuc-MIN6 cells (Kalwat, M.A. et al., 2016b). These cells were treated chronically (24 h) with compounds, preincubated in KRBH in the absence of compounds, and then stimulated. This strategy allowed us to probe a specific pharmacological space to enrich for hit compounds with long-lasting effects. In the screen we stimulated the cells using the diazoxide paradigm to drive both the triggering and metabolic amplifying pathways of insulin secretion (Kalwat, M.A. and Cobb, M.H., 2017, Henquin, J.C., 2000). In this paradigm, diazoxide locks the K_ATP_ channel open, preventing Ca^2+^ influx and insulin secretion until extracellular KCl is provided to depolarize the membrane and elicit the triggering pathway. Addition of glucose under these conditions results in the metabolic amplification of insulin secretion without any further increase in Ca^2+^ influx. This approach provided a platform to later assign the actions of hit compounds to triggering and/or amplification and gave maximal dynamic range in the assay. From the screen, we selected natural product fractions that appeared non-toxic after 24 h, but still potently inhibited glucose-stimulated insulin secretion. Among the top hits was fraction SNB-022-1 derived from *Streptomyces anulatus.* In rescreening we found this fraction retained the expected inhibitory activity (**Fig 1A**). Further fractionation, high resolution mass spectrometry analyses and NMR analyses showed the major component of the fraction to be the aureolic acid chromomycin A_2_ (CMA2) in subfractions F and G (**Fig 1B**). The structure of CMA2 is shown next to an aureolic acid family member, mithramycin, for comparison (**Fig 1C**). We determined the EC_50_ of CMA2 in our reporter assay to be 11.8 nM (**Fig 1D**). Acute treatment (2 h) with CMA2 was insufficient to inhibit glucose-stimulated InsGLuc secretion in our reporter β cells (**Fig 1E**), arguing against a rapid mechanism of action. Importantly, we confirmed the inhibitory effects of 24 h treatment with CMA2 on glucose-stimulated insulin secretion from human islets (**Fig 1F**).

**Figure 1.**
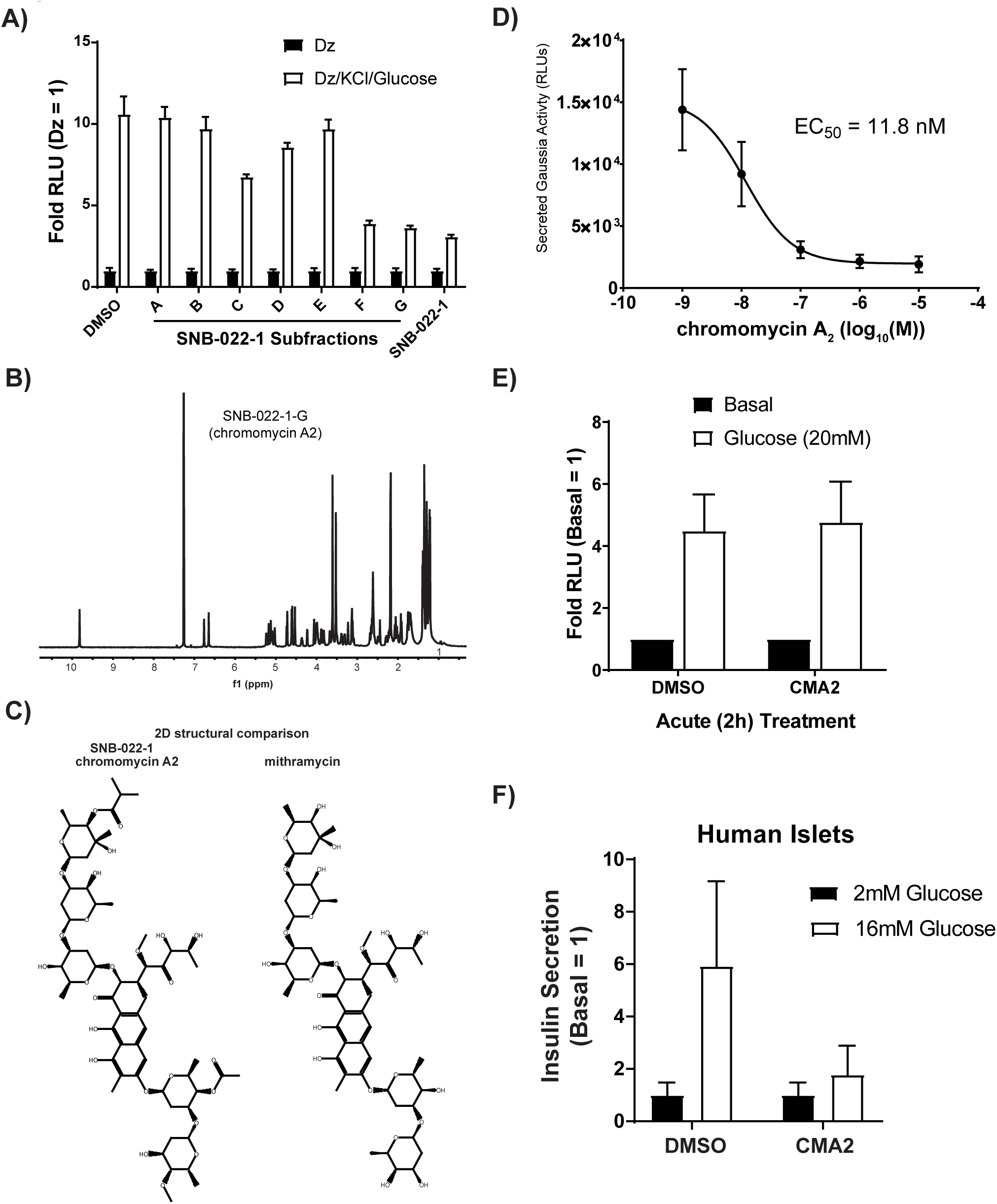
Activity-guided fractionation identifies chromomycin A2 as an insulin secretion inhibitor in human islets. **A**) Natural product subfractions from SNB-022-1 were tested in quadruplicate for inhibitory activity in Insulin-Gaussia Luciferase (Ins-GLuc) MIN6 β cells stimulated with the diazoxide (Dz) paradigm. Cells were treated 24 h with the original SNB-022-1 fraction, subfractions (10 μg/ml) or DMSO (0.1%). Cells were preincubated in glucose-free KRBH for 1 h followed by 1 h in the presence of either Dz (250 μM) or Dz + 40 mM KCl and 20 mM glucose. Secreted luciferase activity is shown. **B**) Nuclear magnetic resonance spectra of the purified compound from fraction SNB-022-1 identified as chromomycin A2. **C**) The spectra in (B) matched chromomycin A2 and the structure is shown. The structure of the closely related mithramycin is shown for comparison. **D**) Ins-GLuc MIN6 cells were treated 24 h with different doses of CMA2 followed by a glucose-stimulated secretion assay and the resulting RLUs were used to determine the effective concentration EC_50_. Data are the mean ± SE (N=4). **E**) Ins-GLuc MIN6 cells were treated with CMA2 (100 nM) starting at the beginning of the 1 h KRBH preincubation and during the 1 h glucose (20 mM) stimulation. Secreted luciferase activity was measured at the end of the assay. Data represent the mean ± SE of three independent experiments performed in triplicate. **F**) Human islets were hand-picked into groups of 50 and treated 24 h with CMA2 at the indicated doses in 500 μl of complete CMRL medium. The islets were then subjected to static glucose-stimulated insulin secretion assays and the secreted and total insulin concentrations were measured. Data are the mean ± SE from three different batches of donor islets.

### Chromomycin A_2_ and mithramycin have differential impacts on triggering and amplifying insulin secretion and gene expression in β cells

Given what is known about mithramycin-DNA binding activity (Hou, C. et al., 2016), the effect of mithramycin on gene expression in glioblastoma models (Singh, D.K. et al., 2017), and the structural similarities between CMA2 and mithramycin (**Fig 1C**), we tested CMA2 and mithramycin in our insulin secretion reporter assays. Interestingly, CMA2 was about 50 times more potent than mithramycin at inhibiting glucose-stimulated secretion after a 24-h incubation (**Fig 2A**). We assessed cell viability after secretion experiments using the Cell Titer Glo assay, which indirectly measures relative ATP concentrations. Interestingly, CMA2-treated cells had an elevated Cell Titer Glo signal while mithramycin-treated cells did not (**Fig 2B**).

**Figure 2.**
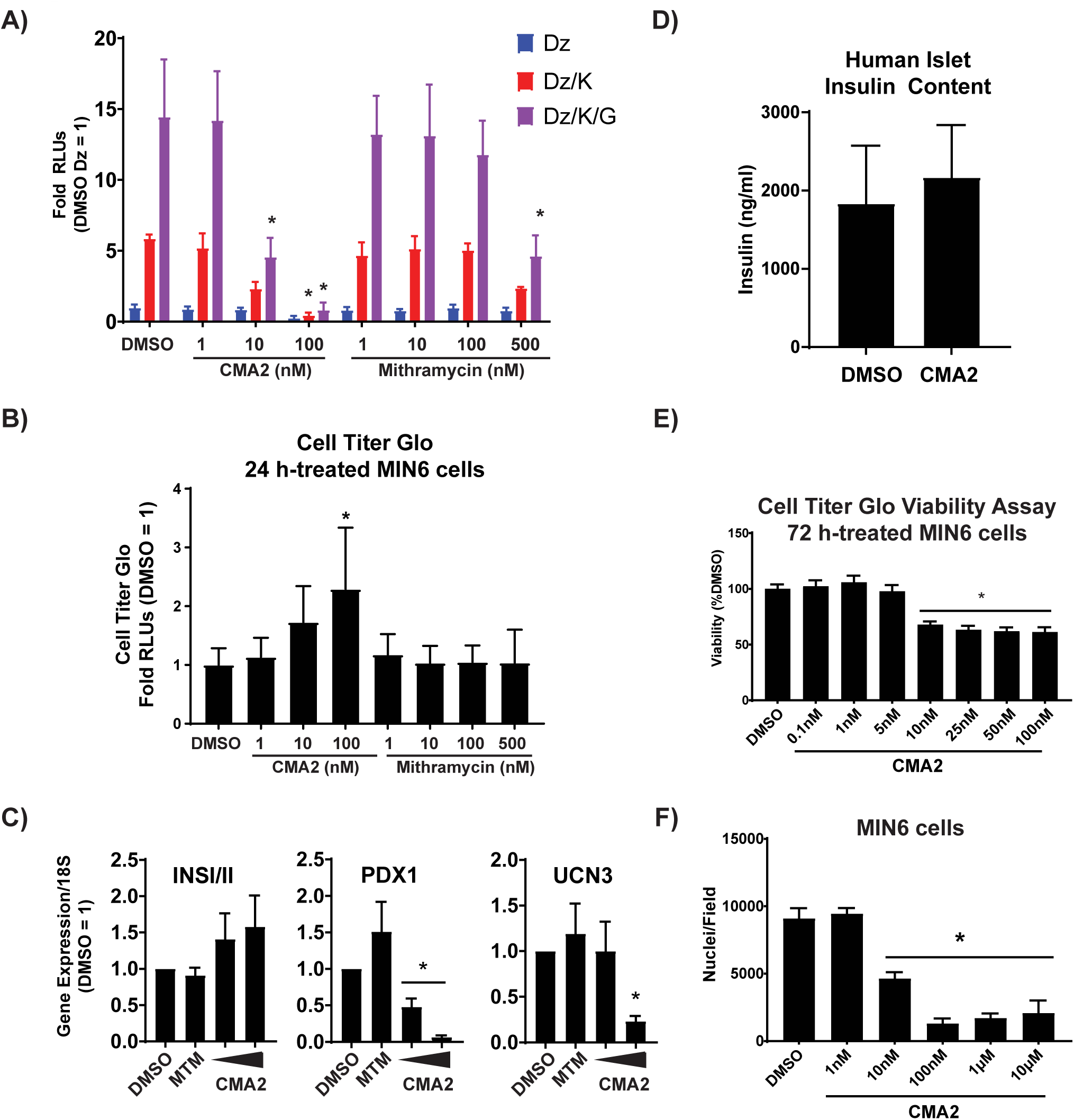
CMA2 has distinct effects on β cells compared to the analog mithramycin. **A**) Ins-GLuc MIN6 cells were treated 24 h with either CMA2 or mithramycin (MTM) at the indicated doses and then secretion was stimulated using the diazoxide (Dz, 250 μM) paradigm as previously. K, 40 mM KCl. G, 20 mM glucose. Secreted luciferase activity was measured and the data are the mean ± SE from three independent experiments. *, P<0.05 by two-way ANOVA. **B**) Lysates of the diazoxide-treated samples in (**A**) were subjected to Cell Titer Glo assays to measure viability. The luciferase activity is shown with respect to DMSO control and the data are the mean ± SE (N=3). **C**) MIN6 cells were treated with CMA2 for 24 h and RNA was isolated for RT-qPCR against INSI/II, PDX1, and UCN3. Data were normalized to 18S rRNA by the 2^^−∆∆Ct^ method and are the mean ± SE (N=3). *,P<0.05 vs DMSO control. **D**) Insulin content was measured in lysates from human islets that were treated 24 h with CMA2 (100 nM) or DMSO (0.1%). Data are the mean ± SE (N=3). **E**) MIN6 cells were treated with the indicated doses of CMA2 for 72 h and subjected to a Cell Titer Glo assay to measure viability. Data are the mean ± SE normalized to the percent of DMSO (N=3). *, P<0.05 vs DMSO. **F**) The data in (**D**) were confirmed by treating MIN6 cells with CMA2 for 48 h followed by fixation and staining with DAPI. Nuclei counts per well are shown as the mean ± SE (N=3). *,P<0.05 vs DMSO.

Aureolic acid compounds are known to influence transcription, therefore we hypothesized that CMA2 may exert some of its effects on β cells by suppressing the expression of critical genes. To assess the effects on gene expression we treated MIN6 cells with CMA2 or mithramycin for 24 h and extracted RNA for RT-qPCR analysis. CMA2, but not mithramycin, suppressed expression of an important β cell transcription factor, PDX1, at as low as 10 nM (**Fig 2C**). Expression of Urocortin 3 (UCN3), a marker of β cell maturation and paracrine regulation of somatostatin secretion (van der Meulen, T. et al., 2015, van der Meulen, T. et al., 2012), was also suppressed but only at 100 nM CMA2, while insulin gene (INSI/II) expression was unaltered (**Fig 2C**). CMA2 (100 nM, 24 h) also had no effect on insulin content in human islets (**Fig 2D**). While treatment with CMA2 for 24 h increased the Cell Titer Glo signal in MIN6 cells (**Fig 2B**), a longer-term 72-h treatment resulted in reduced viability as determined by the same assay (**Fig 2E**) and reduced cell number determined by counting nuclei (**Fig 2F**).

### Chronic CMA2 treatment interferes with triggering Ca^2+^ influx in MIN6 β cells

We next measured Ca^2+^ influx in MIN6 cells pre-treated for 24 h with 100 nM CMA2. In response to glucose, Ca^2+^ influx was not different between vehicle- and CMA2-treated cells (**Fig 3A**), suggesting the dramatic ablation of glucose-stimulated insulin secretion was not primarily due to a defect in glucose-stimulated Ca^2+^ influx. However, we also tested Ca^2+^ influx under the diazoxide paradigm. Diazoxide and KCl clamped the Ca^2+^ levels as expected, as glucose stimulation did not cause additional changes in intracellular calcium levels (**Fig 3B**, green vs orange and purple vs black). With a 24-h CMA2 pretreatment, we did observe a suppression of the triggering Ca^2+^ influx (**Fig 3B**).

**Figure 3.**
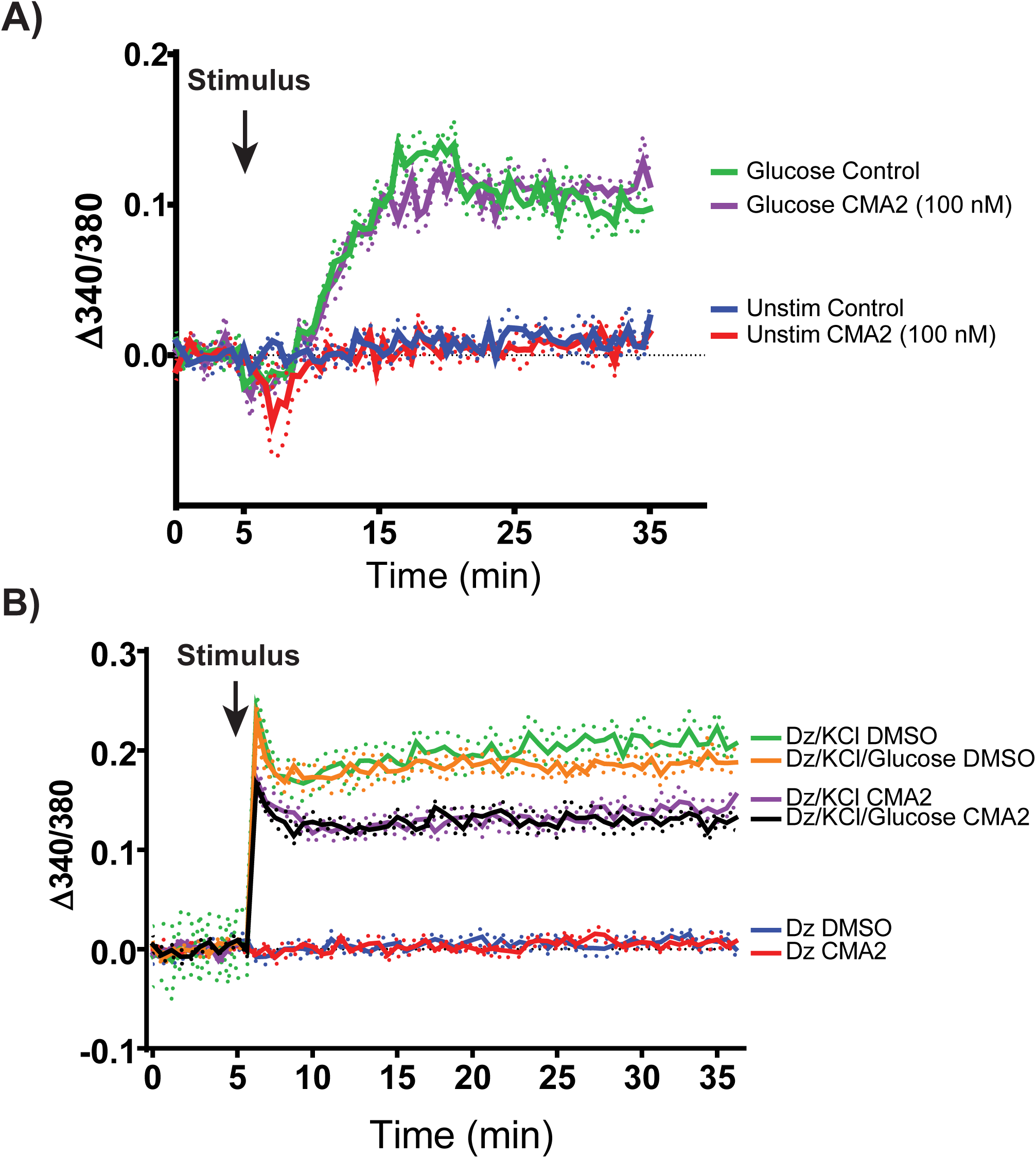
Diazoxide paradigm unmasks effect of CMA2 on triggering Ca^2+^ influx. MIN6 cells were treated 24 h with CMA2 (100 nM) and then prepared for Ca^2+^ influx measurements using Fura-PE3 (5 μM). In (**A**) cells were preincubated in KRBH without glucose for 1 h, loaded with Fura-PE3 for 30 min, followed by a baseline Ca^2+^ measurement for 5 min. 20 mM glucose was added and Fura fluorescence was measured for 30 min. In (**B**), diazoxide (250 μM) was included in the KRBH during the baseline reading and throughout the stimulation protocol. Data are shown as traces representing the mean ± SE (shown as dots) (N=3).

To address whether well-known glucose-mediated β cell signaling pathways are disrupted by CMA2, we tested glucose-induced phosphorylation of ERK1/2 MAP kinase (Khoo, S. et al., 2003) and ribosomal protein S6 (Guerra, M.L. et al., 2017, Moore, C.E. et al., 2009). MIN6 cells were treated with DMSO or CMA2 for 24 h and the following day preincubated in KRBH containing either DMSO or CMA2 to compare effects on cells treated acutely (2 h in CMA2-KRBH), chronically (24 h in CMA2-medium + 2 h in CMA2-KRBH), or chronically with a washout (24 h in CMA2-medium + 2 h in DMSO-KRBH). After the 2 h KRBH incubation, the cells were stimulated with glucose which increased ERK1/2 phosphorylation by 5 min (**Fig 4A**). Chronic CMA2 treatment suppressed the fold increase in ERK1/2 activation at the 5 min time point. By 30 min all CMA2 treatments had suppressed glucose-stimulated ERK1/2 phosphorylation relative to the 0 min time point (**Fig 4A**). With respect to S6 phosphorylation, in control and acute CMA2-treated cells, pS235/6-S6 was induced by 30 min, whereas both chronic CMA2 treatments blunted this event (**Fig 4A**), in agreement with a growth-suppressive effect of CMA2.

**Figure 4.**
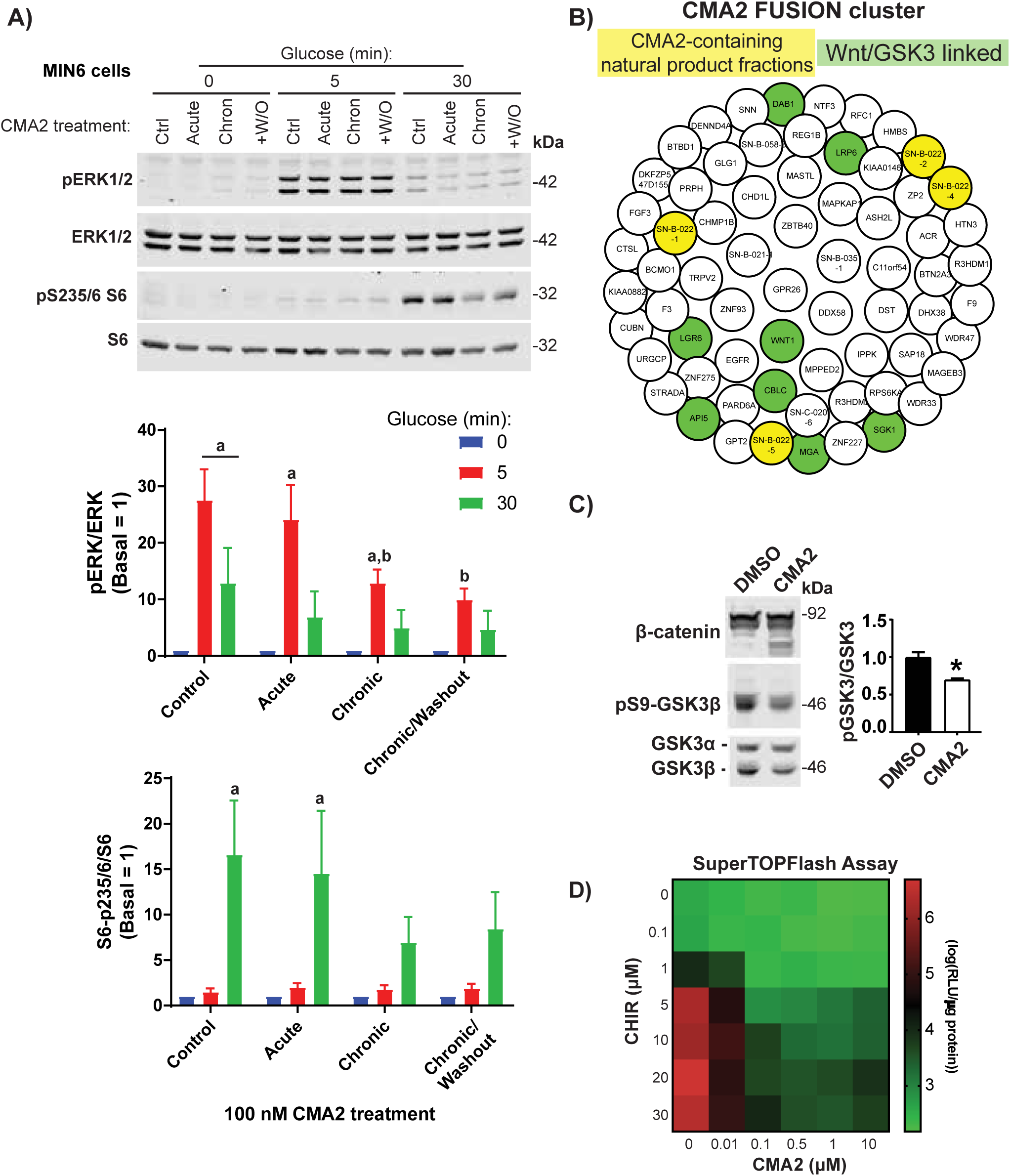
24 h treatment with CMA2 causes dysregulated basal ATP concentrations but does not prevent glucose-stimulated Ca^2+^ influx. **A**) MIN6 cells were treated 24 h with DMSO (0.1%) or CMA2 (100 nM) (chronic) and the next day incubated 2 h in KRBH without glucose, but containing either DMSO or CMA2. Acutely treated cells received CMA2 only during the 2 h KRBH incubation. Washout (+W/O) were cells treated 24 h with CMA2 which received only DMSO during the 2 h KRBH incubation. Cells were then either unstimulated or stimulated with 20 mM glucose for 5 and 30 min and lysed for immunoblotting against pERK1/2, ERK1/2 and pS235/6-S6, and S6. Bar graphs of quantified immunoblots are shown as the mean ± SE (N=3). Using two-way ANOVA: a, P<0.05 vs 0 min glucose. b, P<0.05 vs control, n.s., not significant vs respective glucose 0 min. **B**) FUSION cluster of CMA2 and implicated genes. CMA2-containing natural product fractions are highlighted in yellow and genes linked to the Wnt pathway are highlighted in green. **C**) MIN6 cells treated 24 h with CMA2 (100 nM) or DMSO (0.1%) showed increased β-catenin degradation bands as well as decreased GSK3β Ser9 phosphorylation, indicating increased GSKβ activity. Bar graph shows GSK3β Ser9 phosphorylation as fold of total GSK3β normalized to DMSO control. Data are the mean ± SE (N=3). *, P<0.05. **D**) SuperTOPFlash assay in stable HEK293 cells treated 24 h with different dose combinations of CHIR99021 and CMA2. Displayed data are in log_10_ format. (N=3).

### Chromomycin A_2_ disrupts GSK3/β-catenin pathway signaling in MIN6 β cells

To discern the signaling pathways impacted by CMA2, we used a database developed on campus called FUSION (McMillan, E.A. et al., 2018, Potts, M.B. et al., 2015). FUSION works by measuring genetic perturbation signatures in response to whole genome siRNA, synthetic and natural product screening (McMillan, E.A. et al., 2018, Das, B. et al., 2018, Vaden, R.M. et al., 2017, Potts, M.B. et al., 2015, Potts, M.B. et al., 2013). This approach allows small molecules and natural products to be linked to genes that may be implicated in their mechanisms of action. We applied FUSION to CMA2 and found that suppression of genes connected to Wnt signaling and glycogen synthase kinase 3β (GSK3β) cluster with CMA2. The resulting network allowed us to query what genes may be involved in the actions of CMA2. Through FUSION, we found that CMA2-containing natural product fractions clustered with a small set of genes. Using the online tool Enrichr (http://amp.pharm.mssm.edu/Enrichr/) (Kuleshov, M.V. et al., 2016), we determined this cluster contained Wnt/GSK3β-linked genes (DAB1, LRP6, WNT1, LGR6, API5, CBLC, MGA, SGK1) (**Fig 4B**). In agreement with this finding, chromomycin compounds were previously linked to Wnt pathway inhibitory activity (Toume, K. et al., 2014). When GSK3β is not phosphorylated at serine 9, it is active and can phosphorylate β-catenin, targeting it for proteasomal degradation (Xu, C. et al., 2009). Chronic CMA2 treatment of MIN6 cells also resulted in suppressed GSK3β serine 9 phosphorylation and increased degradation of β-catenin (**Fig 4C**). Wnt pathway activation or small molecule inhibitors of GSK3 (CHIR99021) lead to accumulation of p-catenin, thereby increasing expression of Wnt-responsive genes (Liu, Z. and Habener, J.F., 2010). We observed similar CMA2-mediated inhibition in the β-catenin reporter Super TOPFlash assay (Veeman, M.T. et al., 2003) in HEK293 cells, even in the face of high doses of CHIR99021, which activates reporter expression (**Fig 4D**).

## DISCUSSION

In this study we identified CMA2 as a modulator of β cell function that acts through at least three mechanisms: suppression of gene expression, interference of GSK3/p-catenin signaling, and partial inhibition of Ca^2+^ influx. Although we uncovered certain facets of CMA2 actions, we do not yet know the extent to which each of these mechanisms contribute to the inhibition of insulin secretion; or if there are additional affected mechanisms. Given the potential for aureolic acids to modulate the expression of a wide swath of the transcriptome, future studies will require in-depth gene expression analysis to determine the β cell-specific genes which are disrupted in response to CMA2. Additionally, protein targets of CMA2 or mithramycin have yet to be identified. Elucidation of these targets may explain the differential activity of CMA2 and discovery of the genes and pathways involved in its mechanisms of action could provide in-roads to pharmacologically stabilizing β cell function in a variety of disease settings.

### Inhibitors of β cell function are useful for understanding disease and pathway discovery

It is common to focus on activators of insulin secretion in searching for therapeutics; however, inhibitors are a relatively untapped resource for understanding β cell function. Determining the target genes and pathways of inhibitors of β-cell function, including CMA2, has the potential to elucidate factors that are important for insulin expression, processing and secretion. We identified CMA2 as a Wnt pathway inhibitor in β cells, and the requirements for the Wnt pathway in β-cell development and function are well known (Liu, Z. and Habener, J.F., 2010, Sorrenson, B. et al., 2016, Welters, H.J. and Kulkarni, R.N., 2008, Bader, E. et al., 2016). Therefore, new chemical tools to interrogate this pathway are of immediate relevance to the field of diabetes and islet biology. The uses of aureolic acid natural products in applications such as chemotherapy warrant consideration of side effects on β cells or other sensitive tissues. At the same time, identifying targetable pathways to suppress insulin secretion is also a potentially valuable therapeutic approach for diseases such as polycystic ovary syndrome, insulinomas, and persistent hyperinsulinemic hypoglycemia of infancy. In some cases these disorders are treatable with diazoxide, but there are individuals who are unresponsive to therapy (Snider, K.E. et al., 2013). Insulin secretion inhibitors such as diazoxide, in conjunction with exogenous insulin therapy, have also been used to ‘rest’ β cells, thereby prolonging endogenous functional β cell mass (van Boekel, G. et al., 2008, Brown, R.J. and Rother, K.I., 2008), as well as preserving isolated human islet function prior to transplant (Nijhoff, M. et al., 2018). Therefore, identifying and segregating CMA2 target pathways will be useful for future development of chromomycin analogs.

### Cytoxicity-dependent and -independent impacts of CMA2 on β cell function

We observed that CMA2 at high doses and longer durations (48-72 h) of treatment also exhibited cytotoxic or cytostatic effects. This is likely due in part to the suppression of gene expression and blunting of pro-survival β-catenin activity. Consistent with the impact of CMA2 on GSK3/β-catenin signaling in β cells, another small molecule Wnt signaling inhibitor and GSK3 activator, pyrvinium, led to dramatic suppression of glucose-stimulated insulin secretion (Sorrenson, B. et al., 2016). Additionally, CMA2 was identified to inhibit DNA gyrase *in vitro* at concentrations above 50 nM (Simon, H. et al., 1994); however, its Wnt pathway inhibitory activity is observed in cells at doses below 5 nM, suggesting dose-dependent target selectivity. This is in-line with our observations of an 11.8 nM EC_50_ for inhibition of glucose-stimulated insulin secretion by CMA2 and the ability of 10 nM CMA2 to suppress CHIR-induced β-catenin activity. While CMA2 impacts cell growth, its suppression of insulin secretion may have independent mechanisms. For example, another natural product with growth-suppressive activities, everolimus, was recently shown to suppress insulin secretion independent of its growth inhibition effects (Suzuki, L. et al., 2018).

In many of the experiments in which cells were treated chronically (24 h) with CMA2, the medium and compound were washed off with KRBH prior to the 1-2 h preincubation period and stimulation. This suggests a lasting impact of CMA2 on β cells, perhaps through effects on transcription or protein stability, or that CMA2 remains inside cells even after a washout. Because 100 nM CMA2 pre-treatment almost completely blocked secretion in both triggering and amplifying conditions (**Fig 2A**), we surmise that the relative reduction in KCl-stimulated Ca^2+^ influx likely contributes to, but does not entirely account for the inhibition of secretion (**Fig 3B)**. Our findings that 1) CMA2 treatment did not suppress insulin expression or insulin content, and 2) CMA2 did not blunt Ca^2+^ influx in response glucose stimulation alone, suggest that CMA2 disrupts factors required downstream of Ca^2+^ for insulin secretory granule transport, docking, or exocytosis. Further experiments focused on granule trafficking will be required to determine the specific components affected by CMA2.

### CMA2 and mithramycin elicit divergent phenotypes in β cells

We investigated whether the structurally similar CMA2 relative mithramycin had a similar effect on secretion, but to our surprise, as much as 50 times more mithramycin was required to have the same impact on secretion as 10 nM CMA2. While 100 nM CMA2 increased relative ATP concentrations after 24 h, 500 nM mithramycin failed to have an impact, indicating other mechanistic differences between these two products. Mithramycin was shown to have tumor selective properties, inhibiting the function of a transcriptional trifecta (Sox2, Zeb1, Olig2) to inhibit growth of glioblastoma while apparently sparing normal tissue (Singh, D.K. et al., 2017). Given the recent work on mithramycin and less toxic isoforms (Osgood, C.L. et al., 2016, Singh, D.K. et al., 2017), and the high similarity between the mithramycin and CMA2 structures, modified forms of CMA2 may prove useful as disease therapeutics or chemical probes for research. Over 30 years ago the differences in potency of the aureolic acid family compounds, including mithramycin and CMA2, were suggested to be a result of differential uptake of the compounds into mammalian cells (Gupta, R.S., 1982). CMA2 and mithramycin were also shown to recognize slightly different C/G-rich DNA tracts (Mansilla, S. et al., 2010). This may be accounted for by small differences in the side chains of CMA2 and mithramycin (Barcelo, F. et al., 2010). While the mechanistic explanation for the differences between CMA2 and mithramycin in β cells are still unknown, it remains possible that in our assays β cells are more sensitive to CMA2 compared to mithramycin due to differential uptake. However, in glioblastoma models 50 nM mithramycin showed substantial effects on gene expression (Singh, D.K. et al., 2017), suggesting 50-100 nM mithramycin is more than sufficient to elicit effects in other mammalian cells. The greater potency of CMA2 suggests nuanced mechanisms that are likely to involve signaling and gene expression critical for insulin secretion. Further clarification of these mechanisms may uncover new targets for diabetes treatment and increase understanding of β cell function.

## Acknowledgments

We would like to thank current and former members of the Cobb lab for helpful discussions. Mithramycin was a generous gift from Ralf Kittler (Eugene McDermott Center for Human Growth & Development). M50 Super 8x TOPFlash was a gift from Randall Moon (Addgene plasmid #12456). Early phases of this work were supported by a National Institutes of Health Ruth L. Kirschstein National Research Service Award DK100113 (M.A.K.) and R01 DK55310 (M.H.C.). Other support was provided through U41 AT008718 (J.B.M.) and R37 DK34128 (M.H.C.). We are grateful to the Welch Foundation for funding portions of this project (I1689 J.B.M., I1243 M.H.C.).

